# Phagocytosis of SARS-CoV-2 Nucleocapsid- and RNA-Containing Immune Complexes Drives Inflammatory Cytokine Production and Endothelial Dysfunction

**DOI:** 10.64898/2026.04.09.717495

**Authors:** Johannes S. Gach, Daniel Kim, Khoa Vu, Michaela Marshall, Christopher Cachay, Aminah A. Rangwala, Gius Kerster, Delia Tifrea, Eric Pearlman, Matthew Zahn, Marit van Gils, Christopher C. W. Hughes, Donald N. Forthal

## Abstract

The aberrant inflammation that characterizes severe COVID-19 is incompletely understood. Given the persistence of SARS-Cov-2 RNA and nucleocapsid protein (N) and the presence of anti-N antibody during the course of severe infection, we investigated the role of RNA-containing immune complexes (ICs) in driving inflammation. We found that ICs consisting of SARS-CoV-2 RNA, N, and anti-N IgG1 are internalized by primary human monocytes and stimulate inflammatory cytokines and chemokines in a manner dependent on Fcγ receptors and partially dependent on toll-like receptor-8. In addition, the inflammatory response induced in monocytes by RNA-containing ICs caused endothelial dysfunction in vascularized micro-organs. Using nasopharyngeal samples from SARS-CoV-2-infected individuals, SARS-CoV-2 RNA and N were captured by anti-N monoclonal antibody in the absence of lysing reagents, indicating that SARS-CoV-2 RNA and N complexes are present outside of virions and cells. Finally, we found that during an early wave of COVID-19, higher anti-N IgG:IgM ratios were associated with more severe clinical outcomes, consistent with a role for inflammatory, IgG-mediated phagocytic clearance of nucleic acid-containing ICs in SARS-CoV-2 pathogenesis, perhaps mitigated by non-inflammatory, IgM-mediated clearance. We conclude that RNA-containing ICs may play a role in the pathogenesis of severe COVID-19. Since all pathogenic viruses encode nucleic acid-binding proteins, such as N, and these proteins often elicit an antibody response, inflammatory clearance of nucleic acid-containing ICs may also contribute to disease severity in other viral infections.

## INTRODUCTION

Most cases of COVID-19 are mild and self-limited. However, severe infections with the causative agent, SARS-CoV-2, can result in acute respiratory distress syndrome, widespread endothelial damage and thrombosis, multi-organ failure, and death^1–4^. Aberrant inflammation, associated with risk factors such as advanced age, male sex, obesity, and certain co-morbid conditions underlies severe outcomes^5–7^. However, the fundamental driver of the aberrant inflammation associated with severe disease remains elusive.

In most patients, infectious virus is no longer measurable after about 8 days of infection, presumably due to viral clearance by innate and adaptive effector mechanisms^8^. However, SARS-CoV-2 RNA frequently persists long after viable virus can be isolated, and more severe COVID-19 cases tend to have greater levels and a longer duration of viral RNA in respiratory secretions^9^. Thus, it is plausible that severe disease is linked to the persistence of viral RNA rather than to actively replicating virus and that viral RNA is directly involved in pathogenesis.

The SARS-CoV-2 nucleocapsid protein (N) is the most abundant protein encoded by the virus^10^. Among other functions, N plays a critical structural role by binding viral RNA^11,12^. Moreover, N is highly immunogenic, eliciting in the second week of infection a transient IgM response and a more enduring IgG response^13,14^.

We postulated that RNA bound to N forms immune complexes (ICs) with anti-N IgG and that phagocytosis of such tripartite ICs is highly inflammatory. The internalization of nucleic acid-containing ICs, like those formed with anti-double stranded DNA (dsDNA) or anti-Smith IgG in systemic lupus erythematosus (SLE), is known to stimulate the production of cytokines and chemokines in an Fcγ receptor (FcγR)- and toll-like receptor (TLR)-dependent manner and may be involved in the pathogenesis of autoimmune disorders^15–18^. By analogy, the internalization of ICs containing SARS-CoV-2 N, RNA, and anti-N IgG by phagocytes such as monocytes, macrophages, or dendritic cells might similarly result in inflammation in the setting of SARS-CoV-2 infection.

In this report, we demonstrate that monocyte phagocytosis of nucleic acid-containing ICs composed of N, SARS-CoV-2 RNA, and anti-N monoclonal IgG1 antibodies triggers a response that includes the secretion of inflammatory cytokines and chemokines. Moreover, using a vascularized micro-organ (VMO) model, we show that in the presence of monocytes, the ICs induce significant endothelial cell leakiness, providing a mechanistic link between IC clearance and vascular dysfunction. Finally, consistent with an IgG-mediated pathological response, we provide clinical evidence that patients with severe COVID-19 exhibit higher anti-N IgG:IgM ratios than those with milder disease.

## MATERIALS AND METHODS

### Ethics statement

Peripheral blood from anonymous healthy donors was obtained for this research through the University of California, Irvine (UCI) Normal Blood Donors Program (HS# 2014-1709). Informed, written consent was obtained from all participants in accordance with the UCI Institutional Review Board.

### Reagents

Plasmids encoding the SARS-CoV-2 N-specific monoclonal antibody (mAb) COVA 103-C12 and spike (S)-specific mAb COVA1-18 were provided by Dr. Marit van Gils (University of Amsterdam). Additional N-specific antibody plasmids were supplied by Dr. Patrick C. Wilson (University of Chicago). The SARS-CoV-2 S expression plasmid (pPPI4 SARS-CoV-2 S NFL 2P Foldon-His) was a gift from Dr. Ilja Bontjer and Dr. Rogier W. Sanders (University of Amsterdam). RNA-encoding plasmids (pJET1.2_FS and pUC57-2019-N) were provided by Dr. Amy S. Gladfelter (Duke University). The vector pLVX-EF1α-SARS-CoV-2-N-2xStrep-IRES-Puro (NR-52973) containing the USA-WA1/2020 N gene was obtained through BEI Resources, NIAID, NIH.

### Antibody mutagenesis

The COVA 103-C12 heavy chain plasmid served as a template to introduce the FcγR-reducing L234A and L235A mutations (LALA variant) and a subsequent P329G mutation (LALA-PG variant) into the Fc region^19^. All amino acid substitutions were introduced via site-directed mutagenesis using the QuikChange XL Mutagenesis Kit (Agilent Technologies, Santa Clara, CA) according to the manufacturer’s instructions. The following primer pairs were utilized: LALA (Forward: 5′-accgtgcccagcacctgaagccgcggggggaccg-3′, Reverse: 5′-cggtccccccgcggcttcaggtgctgggcacggt-3′) and LALA PG (Forward: 5’-tctcgatgggggctccgagggctttgttgg-3’, Reverse: 5’-ccaacaaagccctcggagcccccatcgaga-3’). Mutations were verified via Sanger sequencing at Azenta Life Sciences (Burlington, MA) prior to expression.

### Antibody production and purification

Human embryonic kidney (HEK) 293T cells were cultured in BalanCD HEK293 expression medium (Fujifilm) supplemented with 1× Pen/Strep, 20 mM glutamine and 1× insulin-transferrin-selenium. Cells were transfected with heavy- and light-chain antibody plasmids using polyethylenimine (PEI) at a 1:3 DNA/PEI ratio. Supernatants were harvested 4–5 days post-transfection, cleared by centrifugation (450 × g for 5 min), filtered (0.45 μm), and concentrated using 100-kDa Amicon Ultra-15 centrifugal filters. Antibodies were purified using protein A-Sepharose (Genesee Scientific), eluted with IgG elution buffer (Thermo Scientific) and buffer-exchanged into PBS using 50-kDa Amicon units. Concentrations were determined by assessing UV absorbance at 280 nm using a NanoDrop Spectrophotometer (Thermo Scientific).

### SARS-CoV-2 recombinant protein expression

HEK 293T cells were seeded at 1×10^7^ cells in 143 cm² tissue culture dishes and cultured in serum-free BalanCD HEK293 expression medium (Fujifilm). Cells were transiently transfected with either pLVX-EF1α-SARS-CoV-2-N-2xStrep-IRES-Puro to express the USA-WA1/2020 N-protein or pPPI4 SARS-CoV-2 S NFL 2P Foldon-His for the His-tagged S-protein, using a DNA/PEI ratio of 1:3. At 5–6 days post-transfection, N-expressing cells were harvested and stored at −80°C, while S-containing supernatants were collected, filtered (0.45 μm), and concentrated via 100-kDa Amicon Ultra-15 centrifugal filters (Millipore) for downstream purification.

### CoV-2 N and S purification

For N purification, N-expressing HEK 293T cell pellets were thawed and resuspended in 10 mL of lysis buffer (100 mM Tris/HCl, 150 mM NaCl, 1 mM EDTA, 1% NP-40, pH 8.0) supplemented with 1X Protease Inhibitor Cocktail (Genesee Scientific). The suspension was sonicated using a Cole-Parmer ultrasonic cleaner (four to five 25-s pulses at maximum intensity) and cleared by centrifugation at 12,000 × g for 15 minutes at 4 °C. The cleared lysate was loaded onto a Strep-Tactin XT Superflow column (0.4 mL bed volume, Zymo Research) equilibrated with wash buffer (100 mM Tris/HCl, 150 mM NaCl, 1 mM EDTA, pH 8.0). After 30 minutes incubation at room temperature (RT) on an orbital shaker, the column was washed with 10 mL of wash buffer. N was eluted in 1-mL fractions using elution buffer containing 50 mM biotin.

For S purification, the concentrated HEK 293T supernatant was incubated with 1 mL of Ni-NTA agarose (Genesee Scientific) and equilibrated in binding buffer (50 mM NaPO_4_, 300 mM NaCl, 10 mM imidazole, pH 8.0). Following a 45-minute incubation at RT, the resin was washed with 10 mL of wash buffer (containing 20 mM imidazole). Purified N and S were concentrated and buffer-exchanged into 30 mM HEPES or PBS, respectively, using Amicon Ultra centrifugal filters (30 kDa for N; 50 kDa for S). Protein concentrations were determined by measuring optical density at 280 nm using a NanoDrop spectrophotometer, and purity was assessed via sodium dodecyl sulfate-polyacrylamide gel electrophoresis (SDS-PAGE).

### SDS-PAGE

To assess the purity and molecular weight of recombinant SARS-CoV-2 N and S and antibody variants, SDS-PAGE was performed. Proteins were resolved on 10% or 12% (w/v) polyacrylamide gels using a Mini-PROTEAN Tetra Vertical Electrophoresis Cell (Bio-Rad). Antibody variants were analyzed under both non-reducing and reducing conditions (using 50 mM DTT) to verify assembly and chain purity. Following electrophoresis, proteins were visualized using SimplyBlue^TM^ SafeStain (Thermo Fisher Scientific) according to the manufacturer’s protocol.

### DNA cloning and *in vitro* RNA transcription

Vectors for RNA transcription were kindly provided by Amy Gladfelter, Duke University. The plasmid pUC57-2019-N was used as a template to amplify the SARS-CoV-2 N sequence via PCR using the forward primer 5’-gcactagcggccgccatgtctgataatggacccc-3’ and reverse primer 5’-gagtctagaaagattgccttaggcctgagttgagtc-3’, incorporating *NotI* and *XbaI* restriction sites, respectively^20^. The pJET1.2_FS vector (provided by Amy Gladfelter) was digested with *NotI* and *XbaI* (New England Biolabs (NEB)) to remove the frameshift stimulatory (FS) region and ligated with the digested PCR product using T4 DNA Ligase (NEB)^20^. NEB Stable Competent *E. coli* cells were transformed according to the manufacturer’s protocol. The resulting plasmid, pJET1.2_2019-N, was verified by Sanger sequencing (Azenta).

For *in vitro* RNA synthesis, pJET1.2_2019-N and pJET1.2_FS were linearized with XbaI and gel-purified. RNA was synthesized from 500 ng of DNA template using the HiScribe™ T7 High Yield RNA Synthesis Kit (NEB). Following transcription, DNA templates were removed by incubation with RNase-free DNase I (NEB) for 30 minutes at 37°C. RNA was precipitated with 2.5 M LiCl for 30 minutes at –20°C and pelleted at 16,000 × *g* for 15 minutes at 4°C. Pellets were washed once with ice-cold 70% ethanol, air-dried for 10 minutes, and resuspended in RNase-free water. RNA quality was assessed with a 1% agarose gel containing 1% household bleach as described^21^.

### Electrophoretic mobility shift assay (EMSA)

For EMSA, *in vitro* transcribed SARS-CoV-2 RNA (61 nM; corresponding to 500 ng per 20 µL reaction) was incubated with N (0.61–6.1 µM) to achieve molar ratios ranging from 10:1 to 100:1. Incubations were performed in complexing buffer (30 mM HEPES, 6 mM MgCl_2_, 25 U Promega RNasin® Plus, pH 7.3) for 45 minutes at 37°C. The resulting complexes were resolved on a 1% (w/v) agarose gel containing 1% (v/v) household bleach (Clorox) for RNase inhibition and RNA stabilization. RNA without N served as a negative control to establish a baseline for migration. The RNA was visualized using a Spectroline UV transilluminator (Spectronics Corporation).

### RNase protection assay

*In vitro* transcribed RNA from the N-coding region (500 ng per reaction) was complexed with N at a 1:80 (RNA:N) molar ratio. Complexes were formed in complexing buffer (30 mM HEPES, 6 mM MgCl_2_, pH 7.3) for 45 minutes at 37°C in the absence of RNasin Plus. Following complex formation, varying concentrations (0.001 to 0.1 µg/mL) of RNase A (Thermo Scientific) were added to the reaction mixtures and incubated for an additional 30 minutes at 37°C. RNA in the absence of N served as a control to monitor the efficiency of RNase A-mediated degradation. The resulting RNA fragments were resolved on a 1% (w/v) agarose gel containing 1% (v/v) bleach and visualized as described for the EMSA.

### Isolation and pre-treatment of human monocytes

Monocytes were separated from human peripheral blood mononuclear cells via negative selection using the Miltenyi Biotec Human Classical Monocyte Isolation Kit according to the manufacturer’s instructions. The monocytes were subsequently resuspended in assay medium (RPMI 1640, 10% FBS, 100 U/mL penicillin, 100 µg/mL streptomycin, and 20 mM L-glutamine). Where indicated, the medium was supplemented with either CU-CPT9a (Sigma-Aldrich), a potent and selective inhibitor of toll-like receptor 8 (TLR8), 2.5 μg/mL of goat anti-human CD16A/B, CD32, or CD64 (R&D Systems) or 1 µg/ml of R848. Cells were seeded in 96-well round-bottom tissue culture plates at a density of 1.5×10^5^ cells per well in a volume of 100 µL and maintained at 37°C in a humidified tissue culture incubator (5% CO_2_) until further use. Monocytes were approximately 95% pure.

### Immune complex stimulation assay

To generate ICs, recombinant SARS-CoV-2 N or S was combined with *in vitro* transcribed RNA (61 nM; 500 ng per 20 µL) in complexing buffer at a 1:80 (RNA:protein) molar ratio. The mixture was incubated at 37°C for 30 minutes. IC formation was then induced by adding N-specific or S-specific antibodies at a 1:80 (RNA:antibody) molar ratio, reaching a final reaction volume of 20 µL. After a further 45 minutes at 37°C, the IC formulations were divided equally (10 µL per well) and added to the monocyte-containing tissue culture plate. Following 18 hours of incubation at 37°C, the cell supernatant fluids were collected and stored at -20°C or immediately analyzed for chemokine and cytokine release via ELISA. All assays were performed at least in biological triplicate using monocytes from independent donors.

### Immune complex internalization

To determine if ICs were internalized by monocytes, we used an RT-PCR-based assay similar to that described by Gach, et al^22^. Immune complexes were prepared as described above and incubated with human monocytes in a 96-well round-bottom tissue culture plate. After 90 minutes at 37°C, monocytes were pelleted (1400 rpm for 3 min) to remove the supernatant fluid and washed with 200 µL of wash buffer (1X DPBS containing RNase A (10 µg/mL)). Following a second round of washing, cells were mixed with 150 µL of ice-cold acidic wash buffer (10 mM sodium citrate, 140 mM NaCl, pH 3.0) and spun for 3 minutes at 1400 rpm. Subsequently, the acidic wash solution was removed, and the cells were washed twice with 200 µL of wash buffer. Finally, monocytes were lysed in 100 µL RNA/DNA shield solution and transferred into a fresh plate. Samples were either directly processed for SARS-CoV-2 RNA purification and qPCR or stored at -20°C. qPCR was conducted as described below (Capture assay and qPCR).

### Chemokine and cytokine ELISA

Microtiter plates were coated with 100 ng of capture antibody for CCL4 or IL-1β (R&D Systems) diluted in DPBS and incubated overnight at 4°C. The following day, plates were washed three times with wash buffer (DPBS containing 0.1% Tween-20) and blocked with 5% non-fat dry milk (NFDM) in wash buffer for 1 hour at 37°C. After a single wash, 25 µL of cell stimulation supernatant containing ICs was added to each well and incubated for 90 minutes at 37°C. Plates were then washed four times and incubated with 12.5 ng/well of the corresponding detector antibody for 1 hour at 37°C. Following four additional washes, the plates were incubated with a streptavidin-horseradish peroxidase (HRP) solution for 25 minutes at 37°C. Finally, wells were washed six times, developed using TMB substrate, and the reaction was terminated with H_2_SO_4_. Optical density was measured at 450_nm_ using a BioTek Plate Reader. The results of these experiments are reported as ODs rather than as concentrations, which clearly depict the large changes observed under the conditions studied.

Other human cytokines (IL-1α, CXCL1, CXCL2, TNFα, IL-23, IL-6, S100A8, S100A9, IL-8, and for some experiments, IL-1β) were analyzed according to the R&D Systems manufacturer’s instructions.

### Vascularized micro-organ (VMO) platform fabrication

VMO devices were fabricated, loaded, and maintained following previously described methods^23–25^. Briefly, cord derived-endothelial cells (ECs) were maintained in endothelial growth medium-2 (EGM-2; Lonza) on 0.1% gelatin-coated flasks. Lung fibroblasts (LFs) were maintained in DMEM (Corning) with 10% FBS (Sigma-Adrich).

To generate vascular networks, EPCs (8×10^6^ cells/mL) were mixed with LFs (4×10^6^ cells/mL) and resuspended in 8.0 mg/mL of fibrinogen (Sigma). Thrombin (1 mg/ml; Sigma-Aldrich) was added to the cells) and the mixture was immediately loaded into the VMO tissue chamber (Supplementary Figure 1). The outer channels were coated with 1 mg/mL laminin (Thermo Scientific) to promote EC migration. Pressure heads were established by filling media reservoirs with 350 µL in wells A and B, and 50 µL in wells G and H to maintain flow and pressure gradients (Supplementary Figure 1). Pressure heads were established by filling media reservoirs with 350 µL in wells A and B, and 50 µL in wells G and H, which drove flow along the “artery” (B to H) and “vein” (A to G). Microfluidic resistors set the flow rate from artery to vein (Supplementary Figure 1). After 7 days, which allows a fully developed vascular network, monocytes were mixed with ICs or single components in EGM-2 (total volume of 350 µL) at 4×10^5^ cells/mL and added to well B. Devices were incubated overnight before further tests were conducted.

Vessel integrity and permeability was tested by introducing 70 KDa FITC-dextran into the artery (well B). Fluorescence images were captured every 2.5 minutes for 30 minutes and processed as described^23^.

### Study participants and clinical characteristics

Serum samples were obtained from two different cohorts of patients with COVID-19 infection. The initial cohort (Table 1) was infected in 2020 and included 99 participants (61 males and 38 females) with an average age of 56.0 years (range: 19–95). Due to a single missing entry, the age metric was calculated based on n=98, whereas the full sample (n=99) was retained for all other calculations. Clinical severity within this group was considered mild or moderate (n=73, 73.7%) if the participant was not hospitalized or if hospitalized, did not require care in the intensive care unit or mechanical ventilation. Severe cases (n=26, 26.3%) required ICU care, were mechanically ventilated, and/or died. The mean days to serum collection post symptom onset (DPO) was 14.1 days (SD=17.9). The second cohort (Table 1) was infected between January 2021 and April 2023, when omicron strains of SARS-CoV-2 were prevalent in Orange County, California, and consisted of 79 participants (42 males and 37 females) with a mean age of 60.5 years (range=21–92). Forty-four participants (55.7%) had mild or moderate and 35 (44.3%) had severe disease. DPO was available for 64 of these participants and averaged 8.4 days (SD=4.3; range=3–17). To maintain statistical power, the 15 participants with missing onset data were excluded only from calculations adjusted for DPO.

**Table 1.** Patient characteristics.

| **Cohort 1 (infected in 2020); N = 99** |  | **Cohort 2 (infected in 2021-2023); N = 79** |  |
| --- | --- | --- | --- |
| Mean age, years<br>(range) | 56.0<br>(19 - 95) | Mean age, years<br>(range) | 60.5<br>(21-92) |
| Male, N (%) | 61 (61.6) | Male, N (%) | 43 (53.1) |
| Days post symptom<br>onset, mean (range) | 14.1<br>(1-92) | Days post symptom<br>onset, mean (range) | 8.4<sup>1</sup><br>(3-17) |
| Mild/Moderate<br>Disease, N (%) | 26 (26.3) | Mild/Moderate<br>Disease, N (%) | 44 (55.7) |
| Severe Disease, N (%) | 73 (73.7) | Severe Disease, N<br>(%) | 35 (44.3) |

<sup>1</sup>Data missing from 15 patients.

Nasal swab samples obtained from the UC Irvine Department of Pathology were collected in standard Hardy Diagnostic Transport Medium (Cat #R99) from 20 individuals. SARS-CoV-2 RNA by PCR was positive in 11 of the samples and negative in nine. Among the positive cases, the mean DPO was 3.1 days (SD=1.5; range: 1–5).

### Anti-N IgG and IgM ELISA

ELISA plate wells were coated with 100 ng of recombinant SARS-CoV-2 N or S diluted in DPBS and incubated overnight at 4°C. After three washes, plates were blocked for 1 hour at 37°C. Serial dilutions of patient plasma in blocking buffer were then added and incubated for 1 hour at 37°C. Unbound antibodies were removed by four washes, followed by the addition of a goat anti-human IgG or IgM Fab HRP-labeled conjugate (Jackson ImmunoResearch, 1:1,000 in blocking buffer). After five final washes, bound antibodies were detected and quantified as described above.

### Capture assay and qPCR

Plates were coated with 100 ng/well of either anti-N-specific (COVA 103-C12) or anti-S-specific (COVA1-18) antibodies. Wells were washed with DPBS and blocked (DPBS with 5% NFDM) for 1 hour at 37°C. Nasal swabs, diluted 1:10 in DPBS, were added and incubated for 1 hour at 37°C. To ensure removal of unbound components, wells were washed eight times with DPBS. Captured immune complexes were recovered by adding 100 µL of Zymo Research DNA/RNA Shield, and the mixture was transferred to storage plates for immediate processing or storage at -20°C.

Viral RNA was extracted using a viral RNA isolation kit (Zymo Research) and quantified for N gene-specific copy numbers. The qPCR reactions contained 0.1 µM of each N gene-specific primer (Forward: 5’-GACCCCAAAATCAGCGAAAT-3’; Reverse: 5’-TCTGGTTACTGCCAGTTGAATCTG-3’) and a FAM-labeled probe (FAM-ACCCCGCATTACGTTTGGTGGACC-NFQ-MGB). Amplification was performed using Clara Probe Mix No-ROX (PCR Biosystems) on a Rotor-Gene 6000 (QIAGEN) with the following cycling conditions: 50°C for 30 minutes, 95°C for 15 minutes, followed by 45 cycles of 94°C (15 sec) and 60°C (20 sec). SARS-CoV-2 copy numbers were calculated using a standard curve of N gene-containing plasmids and analyzed in triplicate via Rotor-Gene 6000 series software. All samples were run and analyzed in triplicate.

### Statistics

Data were analyzed using GraphPad Prism 10. For *in vitro* stimulation assays involving multiple conditions, significance was determined using a one-way repeated measures ANOVA with Geisser-Greenhouse correction to account for inter-donor biological variability. Specific between-group comparisons were performed using Šídák’s post-hoc test. Mann-Whitney U or Kruskal-Wallace tests were used for data that were not normally distributed.

Statistical analysis for the RNA internalization assay was performed using a nested one-way ANOVA followed by Dunnett’s post-hoc multiple comparisons test.

For clinical cohort data, continuous variables (e.g., IgG:IgM ratios) were compared between two groups using the Mann-Whitney U test. To identify independent predictors of disease severity while controlling for age, sex, and days post-symptom onset, a multivariable logistic regression model was employed. Results are reported as odds ratios (OR) with 95% confidence intervals (CI). P-values < 0.05 were considered statistically significant.

## RESULTS

### Recombinant SARS-CoV-2 antigens and anti-N IgG variants exhibit the structural and functional integrity required for immune complex assembly

In autoimmune disorders such as SLE, pathogenesis is in part mediated by nucleic acid-containing ICs that are internalized by phagocytes via FcγRs^15–18^. The nucleic acid cargo of the ICs engages TLRs in endosomes resulting in the release of inflammatory cytokines and chemokines. Given that N is an RNA binding protein, we investigated whether ICs composed of anti-N IgG, N, and SARS-CoV-2 RNA would be cleared by a mechanisms similar to autoimmune disorders. For these studies, we expressed and purified the anti-N IgG1 mAbs COVA103-C12, S144-339, S564-98, COVA103-C12 LALA, and COVA103-C12 LALA-PG, the anti-S IgG1 mAb COVA1-18, and the corresponding recombinant N and S proteins. We utilized an HEK293T expression system to ensure post-translational modifications, such as phosphorylation, which affects the RNA-binding function of N^26^. SDS-PAGE analysis confirmed the high homogeneity and purity of both proteins and all antibody preparations (Supplementary Figure 2A-D). Side-by-side analyses confirmed that purified N was biochemically comparable to a commercial SARS-CoV-2 N standard (Sino Biological), exhibiting identical electrophoretic mobility (Supplementary Figure 2A). Similarly, S revealed a band at the expected molecular weight of ∼130 kDa with high purity (Supplementary Figure 2B). The antibody preparations displayed characteristic heavy and light chain bands at approximately 50 kDa and 25 kDa under reducing conditions (Supplementary Figure 2C) and a predominant band with an apparent molecular weight exceeding 180 kDa under non-reducing conditions (Supplementary Figure 2D).

To confirm antibody specificity and functional integrity, we performed a series of ELISAs. All N-specific mAbs demonstrated concentration-dependent binding to N (Supplementary Figure 3A, B). Moreover, the parental antibody COVA103-C12 and its Fc-variants, LALA and LALA-PG, displayed nearly identical high-affinity binding curves, confirming that the Fc-region mutations did not alter antigen recognition (Supplementary Figure 3A). A comparison across the three independent N-specific IgG1 mAbs revealed that S144-339 and S564-98 have similar high affinities, whereas COVA103-C12 exhibited slightly lower apparent affinity (Supplementary Figure 3B). Reciprocal assays confirmed the high specificity of the S-specific antibody COVA1-18 for its target antigen, with no cross-reactivity observed with N, and conversely, no binding of COVA103-C12 to S (Supplementary Figure 3A, C). These results confirm that all reagents were suitable for subsequent experiments evaluating the phagocytosis of nucleic acid-containing ICs.

### N forms a stable complex with RNA and protects the RNA from enzymatic degradation

We next characterized the interaction between SARS-CoV-2 N and its target RNA. Using electrophoretic mobility shift assays (EMSA) and a 1.0 kb synthetic SARS-CoV-2 RNA fragment derived from the N-encoding region of genomic RNA, we found that increasing concentrations of N resulted in a clear shift of the free RNA band to higher molecular weights, indicating the formation of stable N-RNA complexes (Figure 1A). This binding was dose-dependent, and we established that an optimal N:RNA molar ratio of 80:1 resulted in near-complete N saturation.

**Figure 1.**
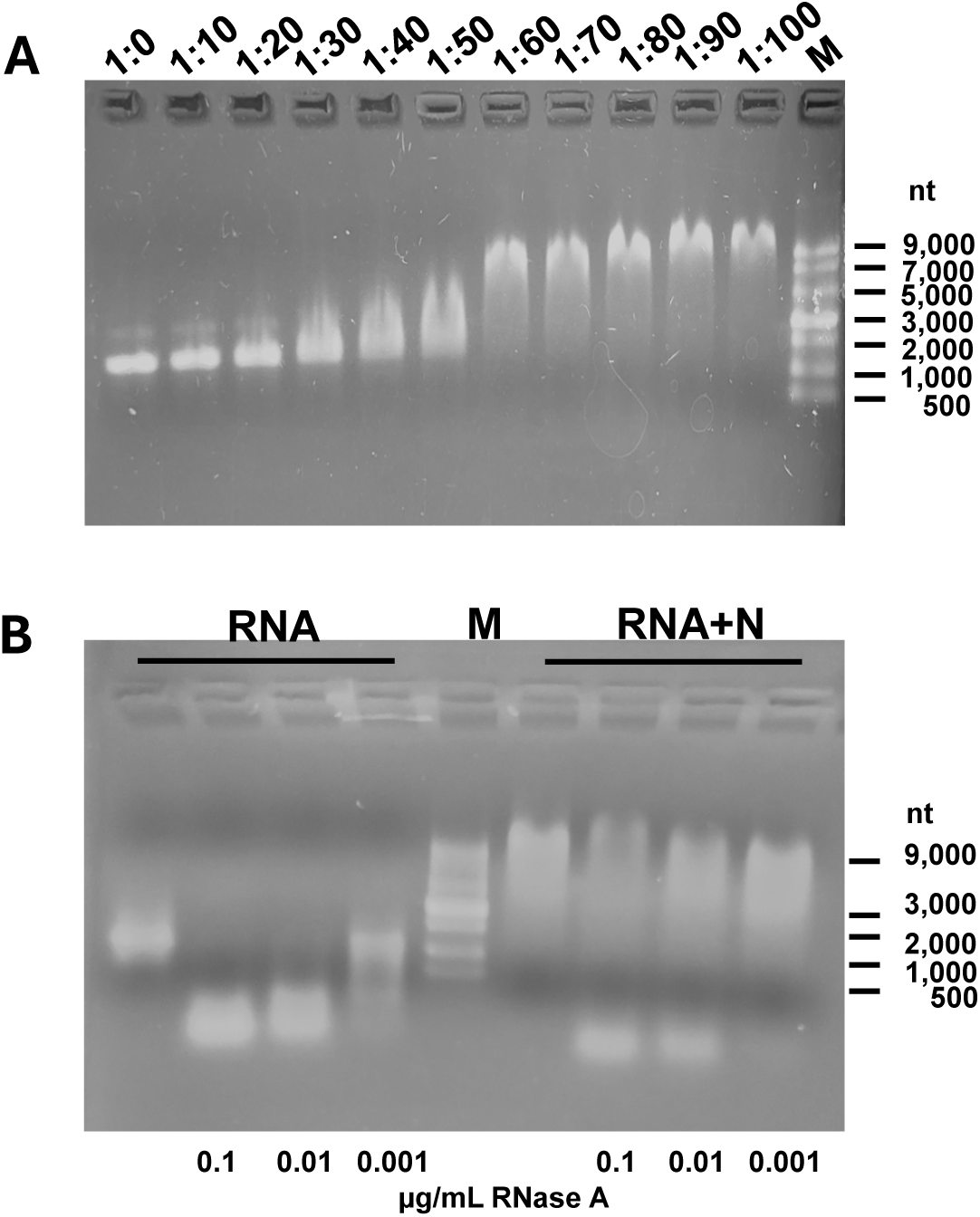
Binding to N protects SARS-CoV-2 RNA from RnaseA degradation. **(A)** *In vitro* transcribed SARS-CoV-2 RNA (from the N-coding region) was incubated with increasing concentrations of purified N to achieve the indicated molar ratios (RNA:N = 1:0 to 1:100). The resulting ribonucleoprotein complexes were resolved on a 1% non-denaturing agarose gel. Lane M contains an ssRNA ladder (sizes in nucleotides, nt, are indicated on the right) used to estimate the size and integrity of the input RNA (∼1200 nt). The progressive reduction in the intensity of the free RNA band and the appearance of a shifted, less mobile band (or smear at lower concentrations) demonstrates a dose-dependent binding interaction. Complete sequestration of the free RNA band is observed at a 1:60 molar ratio and higher. (**B**) *In vitro* transcribed N RNA was incubated with N at a 1:80 molar ratio to form N-RNA complexes (Lanes 5-8). As a control, free RNA was incubated under the same conditions without N (Lanes 1-4). Varying concentrations of RNase A (0.1 µg/mL, lanes 2 and 6; 0.01 µg/mL, lanes 3 and 7; 0.001 µg/mL, lanes 4 and 8) were added to the reactions for 30 minutes at 37 °C. The samples were resolved on a 1% non-denaturing agarose gel. Lane M contains an ssRNA ladder used to confirm the approximate size of the input RNA (∼1200 nt).

We also performed an RNase protection assay to determine if N binding physically shielded the RNA from enzymatic degradation. Free RNA, used as a control, was completely degraded even at low RNase A concentrations (0.1 µg/mL and 0.01 µg/mL; Figure 1B). In contrast, a protected RNA band was visible in the presence of N (80:1 N:RNA ratio) at all tested RNase A concentrations, although some low-molecular-weight degradation products indicated that the protection was not complete.

### Nucleic acid-containing ICs drive pro-inflammatory cytokine production by primary human monocytes

We next determined if the nucleic acid-containing ICs could stimulate cytokine production from monocytes. To generate the ICs, we mixed anti-N IgG1 mAb, N, and SARS-CoV-2 RNA at a fixed molar ratio of 80:80:1, ensuring optimal saturation of RNA-N binding and sufficient IgG density for Fc*γγ*R engagement. The ICs, or controls consisting of the individual IC components or of ICs without nucleic acid, were incubated with primary human monocytes, and after 18 hours, individual cytokines and chemokines were measured in supernatant fluid by ELISA. The results demonstrate that ICs comprising anti-N IgG1, N, and RNA induced secretion of pro-inflammatory mediators associated with COVID-19 or post-COVID-19 sequelae^27–30^, including the cytokines IL-1β, IL-6, IL-8, and TNFα (Figure 2A–D) and the chemokines CXCL1 and CXCL2 (Figure 2E, F). Levels of cytokines and chemokines were often as high as those induced by LPS.

**Figure 2.**
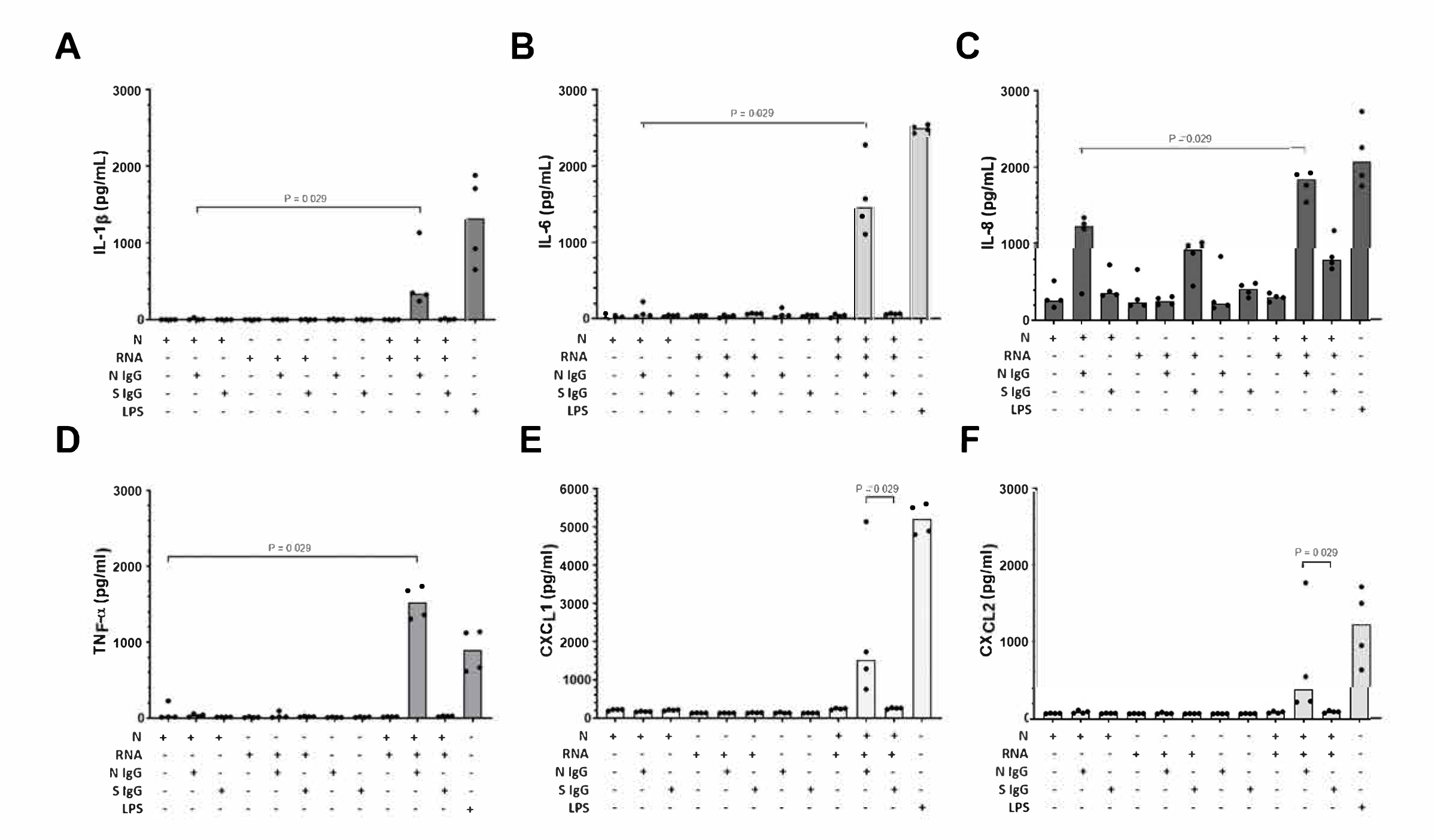
ICs induce monocytes to produce pro-inflammatory cytokines and chemokines associated with severe COVID-19. Primary human monocytes were stimulated for 18 hours with ICs or components of ICs, and the cytokines IL-1β (**A**), IL-6 (**B**), IL-8 (**C**) and TNFα (**D**) and the chemokines CXCL1 (**E**) and CXCL2 (**F**) were measured in supernatant fluid by ELISA. Lipopolysaccharide (LPS) served as a positive control. Data points represent means and standard deviations of 4 independent experiments with different healthy monocyte donors.

We further demonstrated that IL-1β and CCL4 were induced by nucleic acid-containing ICs with two different SARS-CoV-2 RNAs (Supplementary Figure 4A-C) and with three different anti-N IgG1 monoclonal antibodies (Supplementary Figure 4D-E). Chemokine and cytokine production was not induced by ICs made with S or anti-S antibody (Supplemental figure 5). These results collectively demonstrate that ICs containing N, RNA, and anti-N IgG1 specifically trigger monocyte activation and the release of inflammation mediators. Moreover, this inflammatory stimulation by ICs is not restricted to a single sequence of RNA or a single anti-N IgG1.

### Monocyte activation by nucleic-acid containing ICs is dependent on FcγRII

To demonstrate that stimulation of monocytes by nucleic acid-containing ICs made with SARS-CoV-2 components was dependent on FcγRs, we used two Fc-region variants of the anti-N mAb COVA103-C12 containing L234A and L235A (LALA) or LALA plus P329G (LALA-PG) mutations. The Fc-engineered variants, which have reduced binding to FcγRs^19^, retained similar N binding affinities (Supplementary Figure 3A). However, whereas ICs made with the parent IgG1 induced robust secretion of CCL4 and IL-1β, this response was significantly lower with the LALA-PG variant, which does not bind to any of the human FcγRs and does not mediate FcγR-dependent antibody functions (Figure 3A-B)^19^. Thus, cytokine activation by nucleic acid-containing ICs, as expected, is dependent on FcγR binding. Interestingly, the LALA variant (without the PG mutation), which reduces binding to FcγR receptors to a lesser extent than does the LALA-PG variant, generated an inflammatory signal comparable to the unmutated IgG1^19^. Thus, at least for the ICs tested in our assay, there is enough residual Fc-FcγR binding by the LALA variant to result in internalization and activation of monocytes.

**Figure 3.**
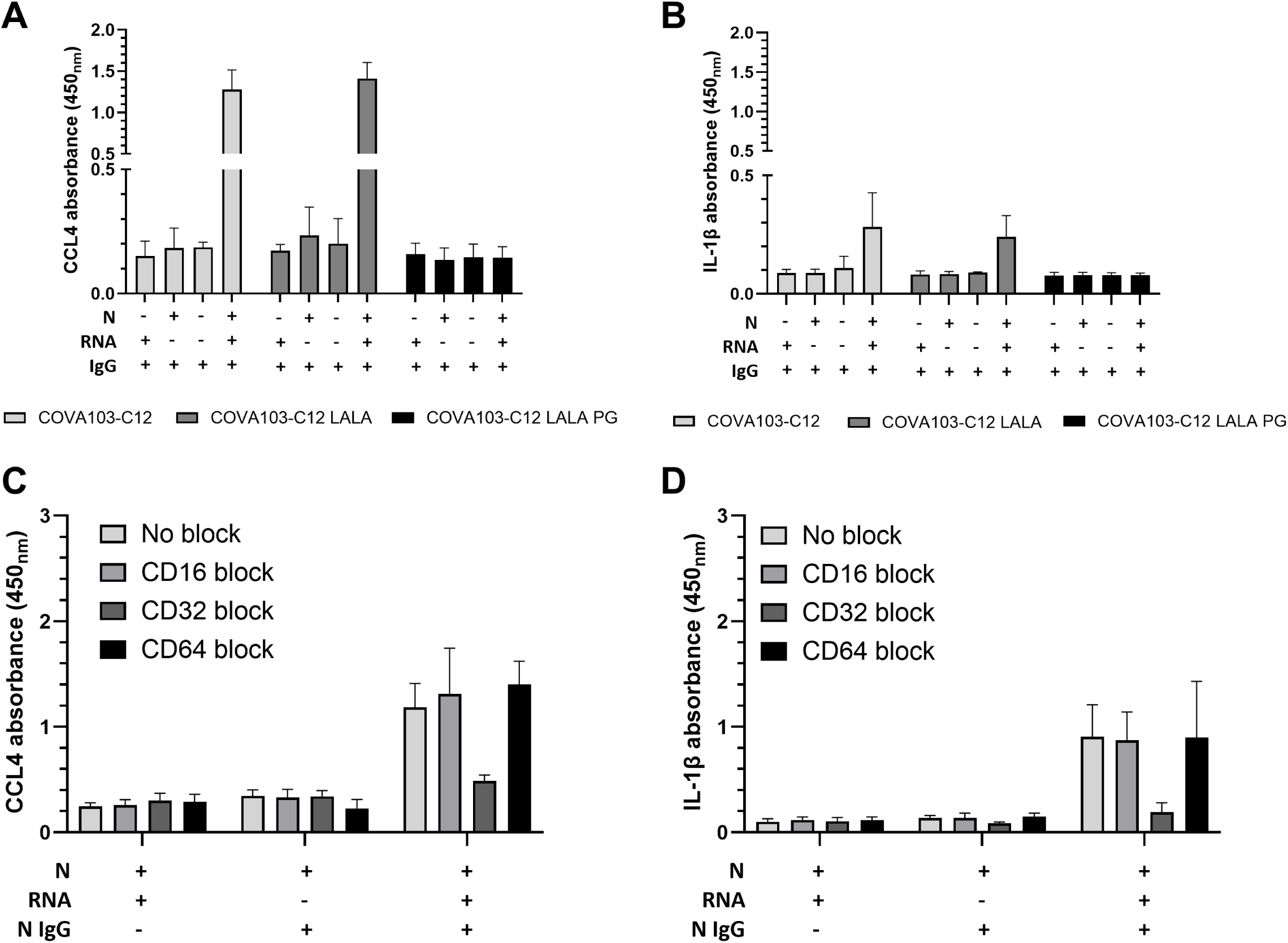
FcγR engagement is required for nucleic acid-containing IC-mediated inflammation. ICs were constructed with wild-type anti-N IgG1 or LALA and LALA-PG Fc variants of the anti-N IgG1 (**A**, **B**). ICs were then incubated with monocytes, and 18 hours later, CCL4 (**A**) and IL-1β (**B**) were measured in supernatant fluid by ELISA. FcγRI (CD64), FcγRII (CD32), and FcγRIII (CD16) were blocked by incubating monocytes with receptor-specific antibodies (2.5 µg/mL) for 1 hour prior to adding ICs (**C**, **D**). 18 hours later, CCL4 (**C**) and IL-1β (**D**) were measured in the supernatant fluid by ELISA. Data represent means and standard deviations of four (**A**, **B**) or three (**C**, **D**) independent experiments with different monocyte donors.

To identify the specific FcγRs that mediate monocyte activation, we stimulated monocytes in the presence of blocking antibodies against FcγRI (CD64), FcγRII (CD32), and FcγRIII (CD16). Anti-CD32 antibody greatly reduced both CCL4 (Figure 3C) and IL-1β (Figure 3D) production by monocytes exposed to nucleic acid-containing ICs. In contrast, blocking CD64 or CD16 had no effect. Thus, engagement of FcγRII is required for inflammatory signaling by monocytes activated by the ICs.

We also determined if nucleic acid-containing ICs were internalized by monocytes. To do so, ICs or controls were added to monocytes for 90 minutes at 37^0^C. After careful washing, the cells were lysed and the amount of SARS-CoV-2 RNA in the lysate was determined by quantitative RT-PCR. SARS-CoV-2 RNA levels in the lysates of monocytes exposed to nucleic acid-containing ICs were about three logs higher than in monocytes exposed to RNA that was not in the form of an IC (Supplementary Figure 6). These results indicate that the ICs are internalized by the monocytes and are consistent with the FcγR engagement required for inflammatory signaling.

### The nucleic acid-containing IC-mediated IL-1β response is dependent on TLR8 signaling

ICs formed with autoantibodies directed against host nucleic acids or ribonucleoproteins engage endosomal TLRs^15,17,18^. Moreover, TLR8 stimulation produces robust and broad cytokine responses in human monocytes^31^. We thus determined the role of TLR8 in IC-induced cytokine production. We found that the TLR8 antagonist CU-CPT9a did not reduce the production of CCL4 induced by nucleic acid-containing ICs (Figure 4A). However, IL-1β production was reduced by approximately 80% in the presence of CU-CPT9a (Figure 4B). The TLR7/8 agonist R848 resulted in increased production of CCL4 and to a lesser extent, IL-1β (Supplementary Figure 7). Thus, CCL4 production can occur directly from endosomal TLR engagement or through FcγR engagement without additional TLR8 signaling^32^. However production of IL-1β, which is primarily regulated through inflammasome activation, does require TLR8 engagement^33^. As expected, CU-CPT9a had no inhibitory effect on the strong inflammatory signal induced by the TLR4 agonist LPS (Figure 4A,B). Thus, monocyte internalization of ICs containing SARS-CoV-2 N, anti-N IgG1, and SARS-CoV-2 RNA is inflammatory and dependent on FcγR and, at least partially, on endosomal TLR-8 engagement.

**Figure 4.**
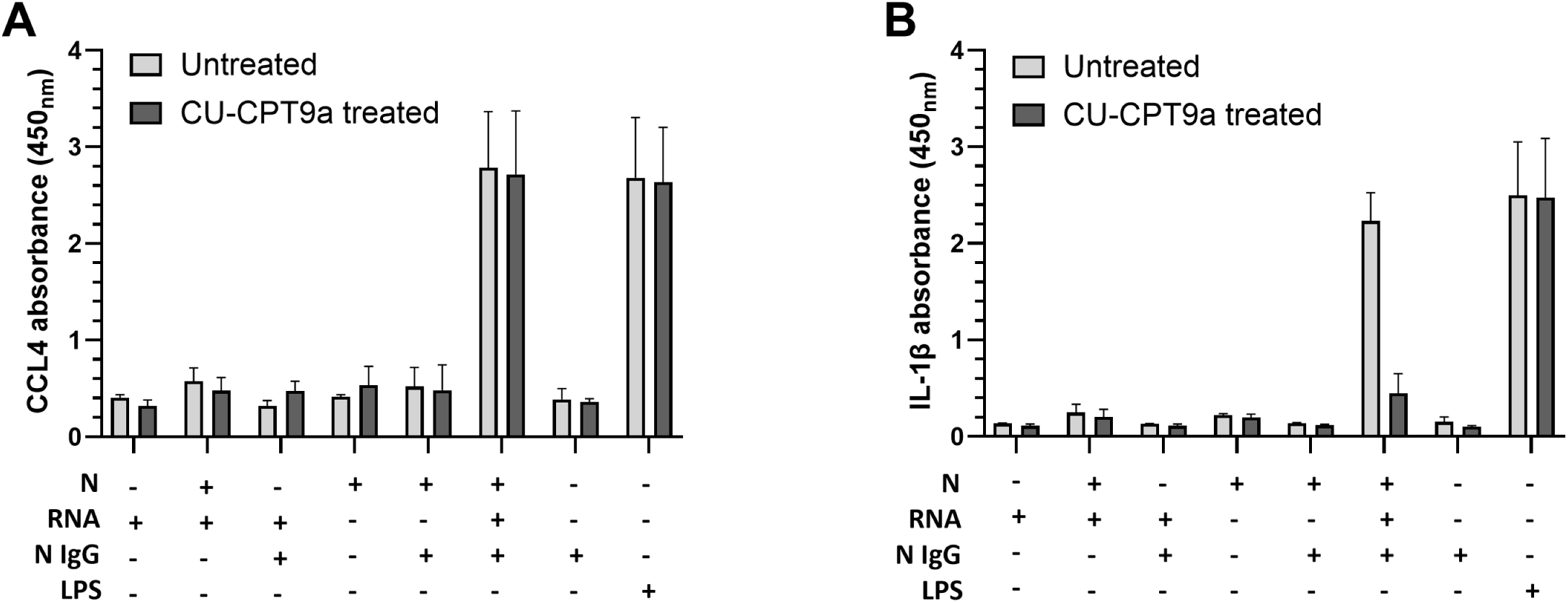
IL-1β but not CCL4 production by monocytes stimulated by nucleic acid-containing ICs depends on TLR8 signaling. CCL4 (**A**) and IL-1β (**B**) was measured by ELISA in the supernatant fluid of monocytes incubated for 18 hours with nucleic acid-containing ICs either in the presence or absence of the TLR8 inhibitor CU-CPT9a. Cells were incubated with 10 µM CU-CPT9a for 2 hours before adding the ICs. Data represent the means and standard deviations of three independent experiments with different monocyte donors.

### Nucleic acid-containing ICs trigger endothelial barrier leakage

Vascularized micro-organs (VMOs) demonstrate key characteristics of human vasculature, such as tight junctions, expression of vascular markers and response to inflammatory stimuli^34,35^. Moreover, they have been used to study the effects of SARS-CoV-2 infection on endothelial cells^4^. Using VMOs, we evaluated the effect of nucleic acid-containing ICs on endothelial function. Endothelial integrity was measured by quantifying extra-luminal fluorescence before and one day after monocytes and ICs were injected into the arteriole of the VMOs (Supplementary Figure 1). We found that the nucleic acid-containing complexes induced a significant increase in leakiness as measured by extra-luminal fluorescence in the presence of monocytes (mean=14.4-fold difference between pre- and post-treatment; p=0.04; Figure 5A, B). As expected, when the poorly FcγR-engaging LALA-PG variant of the anti-N IgG was used, there was little increase in endothelial leakage (Figure 5 C, D). In the absence of monocytes, ICs did not affect vascular leakage (Figure 5E, F). These results point toward a potential role for phagocytosis of nucleic acid-containing ICs in the endothelial dysfunction reported during SARS-CoV-2 infection.

**Figure 5.**
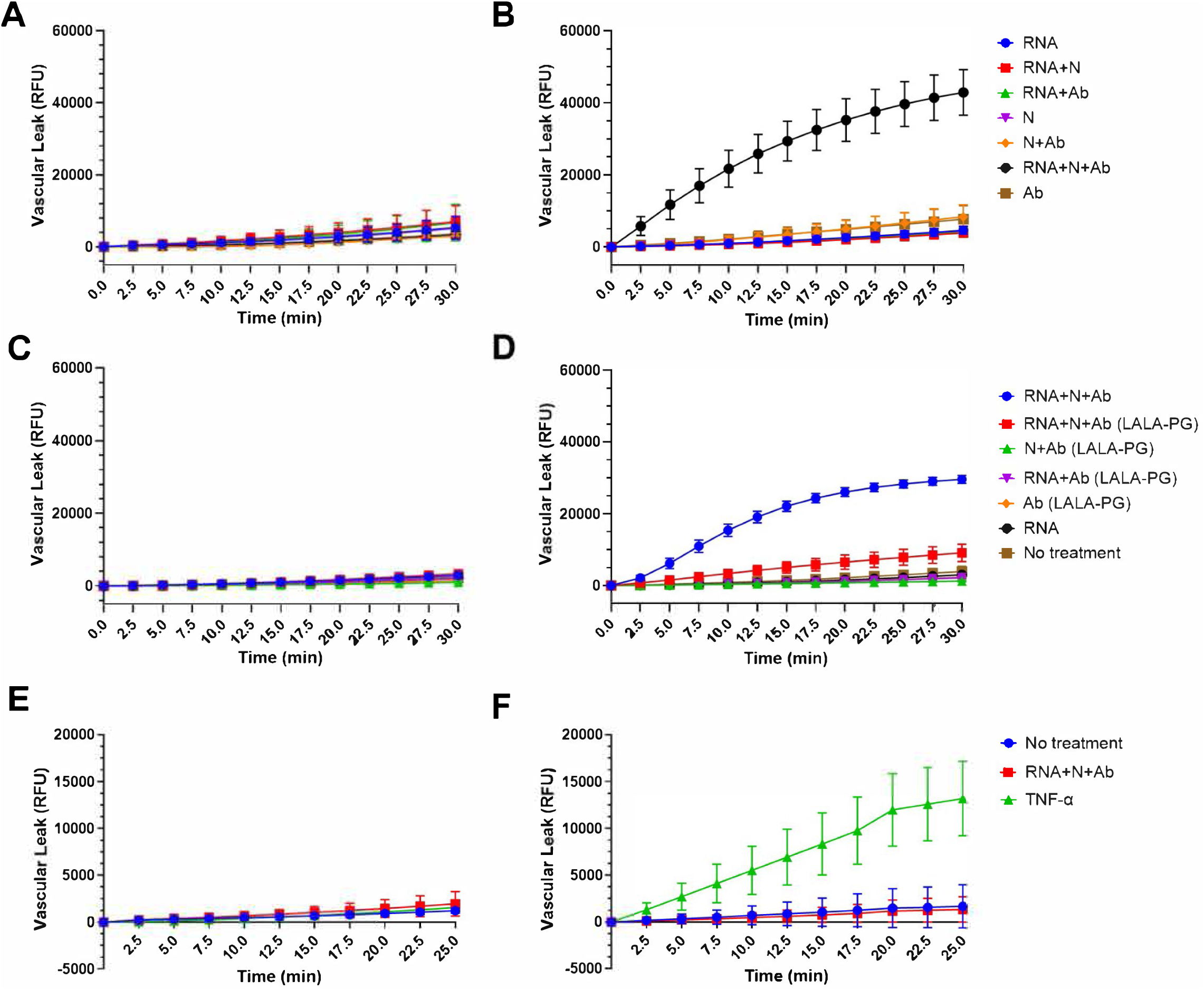
Vascular leakage occurs following exposure of vascular micro-organs (VMOs) to monocytes activated by nucleic acid-containing ICs. Baseline vascular leakage was measured by FITC-dextran extravasation (**A**, **C, E**). 24 hours later, ICs made with the anti-N mAb COVA103-C12 (**B**) or with the LALA-PG variant of COVA103-C12 (**D**) and monocytes were added to the arterioles, and FITC-dextran extravasation was re-measured. Alternatively, COVA103-C12 in the absence of monocytes was added to the VMOs (**F**). TNFα was added as a leak-inducing positive control. Data are recorded as relative fluorescent units (RFUs) and represent means and standard errors of three independent experiments (with different monocyte donors when monocytes were included [**B**, **D]**). Data points for each VMO device pre- and post-treatment are the same color.

### RNA-N complexes naturally occur during COVID-19

If nucleic acid-containing ICs are involved in COVID-19 pathogenesis, there should be evidence that such complexes form naturally during infection. We determined if SARS-CoV-2 RNA bound to N could be detected in nasopharyngeal (NP) swab specimens collected one to five days after the onset of COVID-19 symptoms in 11 patients. Capture was accomplished by coating plates with anti-N or anti-S IgG1 mAbs prior to adding material from SARS-CoV-2-positive or negative NP swabs diluted in PBS. After careful washing, RNA was measured by q-PCR. We did not use lysing compound, thus allowing us to capture both whole virus with anti-S mAb and virion-free N-RNA complexes with anti-N mAb. Anti-N mAb and anti-S mAb captured RNA from 6 and 7 NP samples, respectively (Figure 6A, B). Thus, at least during early stages of infection, N in complex with RNA is detectable in respiratory secretions from patients with COVID-19. We did not attempt to capture ICs containing IgG, since during this early period of infection, we would not expect an adequate anti-N IgG response in the upper respiratory tract.

**Figure 6.**
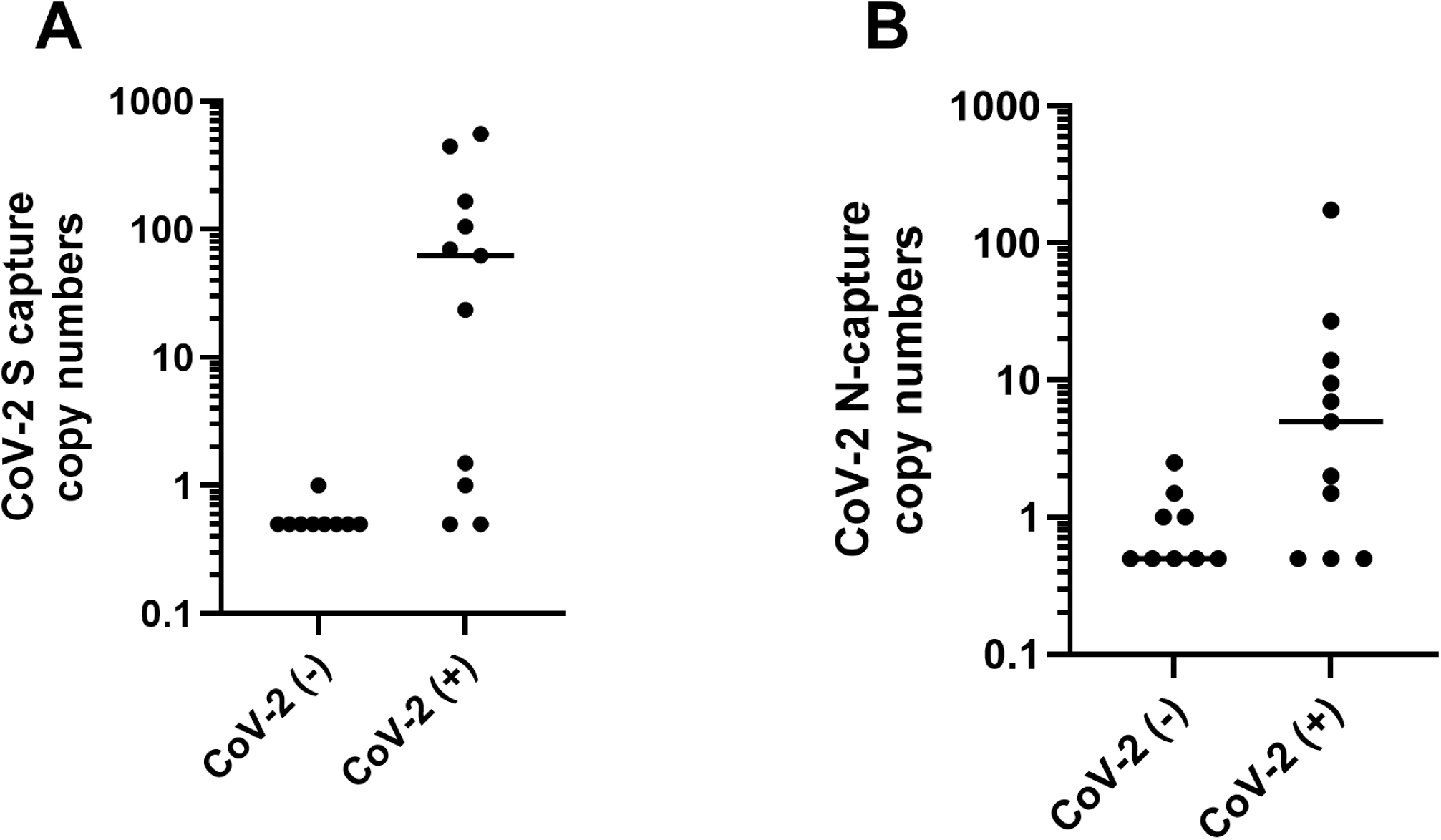
Sars-CoV-2 N-RNA complexes are present in patient nasopharyngeal samples. SARS-CoV-2 RNA was captured by anti-S IgG1 (**A**) or by anti-N IgG1 (**B**) from nasopharyngeal (NP) swab specimens of 11 acutely infected individuals (CoV-2 (+). Nine swabs from individuals without COVID-19 (CoV-2 (-) were included. Each dot represents an individual patient sample. Limit of detection = 5 copies. Horizontal bars indicate the median value for each group.

### Anti-N IgG:IgM ratios were significantly higher in severe compared with mild or moderate disease in the early pandemic period

ICs containing IgG are more likely to be cleared in an inflammatory manner than are ICs containing IgM^36–39^. To further explore the role of nucleic acid-containing ICs in SARS-CoV-2 pathogenesis, we measured serum anti-N IgG and anti-N IgM levels in 99 patients (26 with severe and 73 with mild/moderate disease; see methods for definitions) at a median of 9 days after symptom onset. All patients were infected during the 2020 COVID-19 wave that was due to the original (ancestral) strain. Anti-N IgG levels did not differ between the severe and mild/moderate groups, whereas anti-N IgM trended toward lower levels in the severe group (p=0.06). Since IgG-mediated clearance of ICs is inflammatory and IgM-mediated clearance may be anti-inflammatory^38,40^, we determined if the ratio of the two antibody classes was associated with disease severity. We found that anti-N IgG:IgM ratios were significantly higher in severe than in mild/moderate cases (median 10.2 vs. 5.4 ; p=0.0015; Figure 7A). This association held in a multivariable logistic regression model (p=0.0026) after controlling for age, sex, and days after symptom onset, identifying the anti-N IgG:IgM ratio as an independent predictor of severity (OR=6.7 ; 95% CI: 1.9 to 27.9). By contrast, the anti-S IgG:IgM ratio in the same cohort did not significantly differ between severity groups (median 3.2 vs. 2.0; p=0.12; Figure 7B).

**Figure 7.**
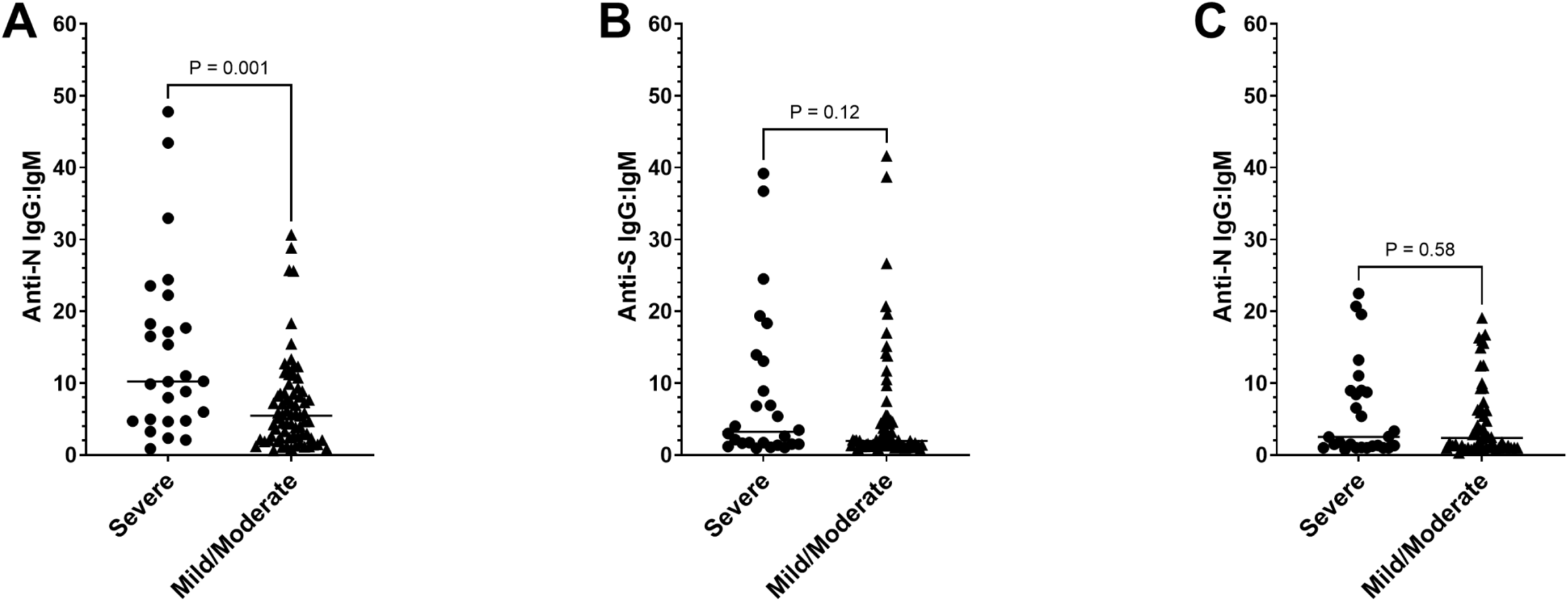
Anti-N IgG:IgM ratios are associated with disease severity during an early wave of COVID-19. Anti-N and anti-S IgG and IgM were measured by ELISA in sera from two cohorts of patients with COVID-19. The anti-IgG:IgM ratio were associated with disease severity in a cohort infected in 2020 (**A**). Anti-S IgG:IgM ratios in the same 2020 cohort did not significantly differ by disease severity (**B**). Anti-N IgG:IgM ratios were not associated with disease severity in a cohort infected in 2021-2023, when omicron SARS-CoV-2 strains were prevalent (**C**). Scatter plots represent individual patient IgG:IgM ratios with horizontal lines indicating median values. Anti-N and anti-S IgG and IgM values were measured twice on each serum, and the average value was used to calculate the IgG:IgM ratio.

To determine if the anti-N IgG:IgM ratio was similarly associated with severity in later waves of the pandemic, we evaluated a second cohort of 76 individuals who were infected between 2021 and 2023; in Orange County, California, ancestral and delta variants predominated in 2021, whereas omicron variants predominated in 2022 and 2023 (unpublished data from Orange County Health Care Agency). Unlike in the earlier cohort, the anti-N IgG:IgM ratio did not differ between those with severe and mild/moderate infection (median = 2.51 and 2.38, respectively; p=0.6; Figure 7C). Thus, the anti-N antibody response may play a role in the pathogenesis of SARS-CoV-2 infection, especially in virulent ancestral strains or in situations where the affected population has little or no prior immunity.

## DISCUSSION

We investigated the potential role of ICs containing N, anti-N IgG, and RNA in the pathogenesis of SARS-CoV-2 infection. We found that these ICs are internalized by phagocytic cells in an inflammatory manner. Moreover, phagocytosis of RNA-containing ICs by monocytes results in endothelial leakage in vascularized micro-organs constructed with human endothelial and stromal cells. We also show that N-RNA complexes are present in the respiratory secretions of patients with COVID-19 and that the RNA in these complexes is protected against degradation by RNase. Finally, during the initial wave of the pandemic, the ratio of inflammatory anti-N IgG antibodies to non-inflammatory IgM antibodies was higher in patients with severe COVID-19 compared to those with mild or moderate infection.

All pathogenic viruses encode a nucleic acid-binding protein, and many of those proteins elicit an antibody response^13,14,41–47^. In the case of SARS-CoV-2, its nucleic acid-binding protein N is the target of an early IgM and IgG response, the latter of which lasts for several months^13,14^. Thus, there is an opportunity for tripartite ICs consisting of anti-N antibody, N, and SARS-CoV RNA to form during infection. These RNA-containing ICs are analogous to those that form between IgG autoantibodies and dsDNA, RNP, or other combinations of protein and nucleic acids during autoimmune disorders such as SLE^15–18^. The nucleic acid-containing ICs in autoimmune disorders are cleared in a highly inflammatory manner that may underlie disease pathogenesis. We now show that exposure of nucleic acid-containing ICs formed with SARS-CoV-2 components results in the production of inflammatory cytokines and chemokines including IL-1β, IL-6, IL-8, TNF-α, CCL4, CXCL1 and CXCL2. These cytokines and chemokines have been associated with COVID-19 severity, pointing toward a potential role for nucleic acid-containing ICs in COVID-19 pathogenesis^27–30^.

In many cases, severe manifestations of COVID-19 become apparent after the first week or two of infection at a time when anti-N IgG is often measurable but infectious virus is infrequently isolated^8,9,13,48^. Viral transcripts and N, however, can persist in tissues or blood for weeks after infection, and higher RNA and N levels are associated with increased severity^9,49–51^. Thus, a pathogenic mechanism that results in aberrant inflammation from phagocytic clearance of lingering nucleic acid-containing ICs could explain the delayed disease severity seen in some patients.

A potential reason for viral RNA persistence is its ability to stably interact with N and resist degradation by host enzymes. Protection from RNase by N has been demonstrated for the coronavirus mouse hepatitis virus^52^. We now show that extracellular SARS-CoV-2 RNA is also protected from RNase degradation when bound to N. Of note, a previous study reported that intracellular genomic RNA is protected by double-membrane cytoplasmic vesicles during viral replication^53^. Our model, however, likely requires that IgG engage RNA—either genomic or subgenomic—outside of cells, and binding to and protection by N may be critical for this to occur. As with ICs made with autoantibodies, the SARS-CoV-2 ICs are internalized via FcγRs.

We show that on primary human monocytes, FcγRII is the primary receptor for the ICs. Moreover, the ICs resulted in the production of inflammatory cytokines, consistent with the cross-linking of the activating receptor FcγRIIa, rather than the inhibitory receptor FcγIIb. As expected, introducing the LALA-PG mutation into the Fc segment of anti-N IgG1, which results in a marked loss of FcγR binding, completely abrogated the production of cytokines. Less expected was our finding that the LALA mutation alone was not sufficient to inhibit cytokine production. This is consistent with some degree of retained LALA Fc-FcγR binding in the absence of further mutations^19^. The apparent high degree of engagement between the IgG1-LALA containing ICs and FcγRII may be explained by the size and multivalency of the nucleic acid-containing ICs.

In the current study, we show that the production of the inflammatory chemokine CCL4, while requiring FcγR engagement, was unaffected by CU-CPT9a inhibition of the endosomal RNA-sensing TLR 8. This suggests that FcγR engagement alone is sufficient or that TLR 7, another endosomal RNA sensor, may be involved^54^. Conversely, IL-1β production was inhibited by pre-incubation with CU-CPT9a. IL-1β production by monocytes is generally dependent on inflammasome activation, and our results, as well as those from other studies, indicate that dual signaling through both FcγRs and TLRs results in inflammasome activation^33,55,56^.

Endothelial dysfunction is a characteristic of severe COVID-19, leading to acute thromboembolic complications and, potentially, to increased cardiovascular events post-COVID-19^3,57–59^. This led us to directly explore the effect of nucleic acid-containing ICs on the vasculature using vascularized micro-organs (VMOs). VMOs are made from human endothelial and stromal cells and allow for physiological flow through arteriole and venule channels. VMOs incubated with nucleic acid-containing ICs resulted in marked endothelial dysfunction as measured by extravasation of FITC-dextran from vessels. The individual components of the ICs or ICs consisting of anti-N IgG1 and N protein without RNA caused little or no vascular leakage. In addition, nucleic acid-containing ICs did not have a direct effect on the endothelium, as demonstrated by lack of increased leakiness in the absence of monocytes. These results, along with our previous data on direct SARS-CoV-2 infection of ECs, provide a potential mechanistic explanation for a key pathological finding during COVID-19^4^. Endothelial dysfunction is also an important phenomenon in autoimmune diseases such as SLE^60^. Recently, the level of anti-Smith antibody (which binds to protein-RNA complexes) was found to positively correlate with levels of circulating markers of endothelial dysfunction^61^. Thus, the inflammatory clearance of nucleic acid-containing ICs could be a generalizable phenomenon in diseases characterized by endothelial dysfunction. Further studies using *in vivo* models will be necessary to determine the impact of IC phagocytosis on vascular dysfunction and its sequelae such as atherosclerosis.

Phagocytic of IgG bound ICs generally results in the production of inflammatory cytokines and chemokines and, as described above, may be pathologically involved in autoimmune disorders. Clearance of IgM bound ICs, however, tends to be non- or anti-inflammatory, possibly due to in part to engagement of IC-C1q complexes with ITIM motif-containing LAIR-1^38,40^. Given the anti-inflammatory nature of IgM and our *in vitro* data demonstrating the inflammatory response to phagocytosis of ICs made of IgG1, N, and RNA, we postulated that a higher anti-N IgG:IgM ratio would be associated with worse COVID-19 outcomes. We found that during an early wave of COVID-19, patients with severe infection, defined as requiring ICU admission, mechanical ventilation or death, had higher anti-N IgG:IgM ratios than patients with mild or moderate disease. No such association was found with anti-S antibodies, which would not be expected to form nucleic acid-containing ICs. The higher ratios were due to a lower anti-N IgM response, rather than higher IgG levels. Elevated IgG responses against S or N have been associated with worse outcomes in several studies, though whether the antibodies themselves are pathogenic or are indicative of a higher antigenic load during severe infection is unclear^62–65^. Our results suggest that anti-N antibody of the IgM subclass could potentially mitigate the pro-inflammatory effects of anti-N IgG antibodies.

We found that the anti-N IgG:IgM ratio was not associated with COVID-19 severity in a cohort of patients who were infected in subsequent waves of infection that largely involved omicron strains. One explanation for this is that strains causing the earlier wave may have been more virulent, and strain-specific antibody responses modified the pathological consequences of IC clearance^66,67^. Relatedly, subsequent waves took place in the setting of some degree of pre-existing anti-SARS-CoV-2 antibodies and T cells from previous infection or vaccination. This prior immunological experience may have lessened the impact of IC clearance on severity. Finally, sample size considerations render it possible that results with either cohort might have been different had larger numbers of subjects been included.

We provide direct evidence that SARS-CoV-2 RNA is bound to N in nasopharyngeal secretions of infected patients. We did not assess non-respiratory samples or have access to samples from individuals with long-term RNA positivity. We also did not determine if the N-RNA complexes were bound by antibody and given the very early stage of infection and the mucosal source of our samples, we did not expect to find ICs bound by IgG. Nonetheless, our results indicate that the foundation for generating nucleic acid-containing ICs, namely the antigen bound to RNA, is present during SARS-CoV-2 infection.

Finally, the presence of anti-N antibody and the persistence of RNA and N in blood, lymphoid tissue, brain, the GI tract or other tissues in some patients with post-acute sequelae of SARS-CoV-2 infection point toward a potential link between ICs and the aberrant inflammatory response and vascular complications observed during long COVID^68–74^.

In summary, we have shown that ICs consisting of anti-N IgG1, N, and SARS-CoV-2 RNA are internalized by monocytes in an inflammatory manner. While analogous to the clearance of nucleic acid-containing ICs in autoimmune disorders, the impact of such ICs from a viral source has not been previously described. Our results suggest a potential pathogenic role for ICs during acute COVID-19 as well as during long COVID. Moreover, since all human viruses encode nucleic acid-binding proteins and many of these proteins elicit antibody responses, nucleic acid-containing ICs could contribute pathogenically to other viral infections.

## ACKNOWLEDGEMENTS

The authors would like to thank Patrick Wilson, University of Chicago, for supplying monoclonal antibody plasmids and Amy Gladfelter, Duke University, for supplying vectors for RNA transcription. We also thank the COSCA study team, in particular Dr. Godelieve J. de Bree, and the participants of the COSCA study.

## Author contributions

JSG, DK, KV, MM, CC, and AAR performed the experiments. GK and MvG provided monoclonal antibodies and plasmids, JSG, EP, CCWH, and DNF designed experiments, interpreted results, and wrote or edited the paper.

## SUPPLEMENTARY FIGURE LEGENDS

**Supplementary Figure 1.**
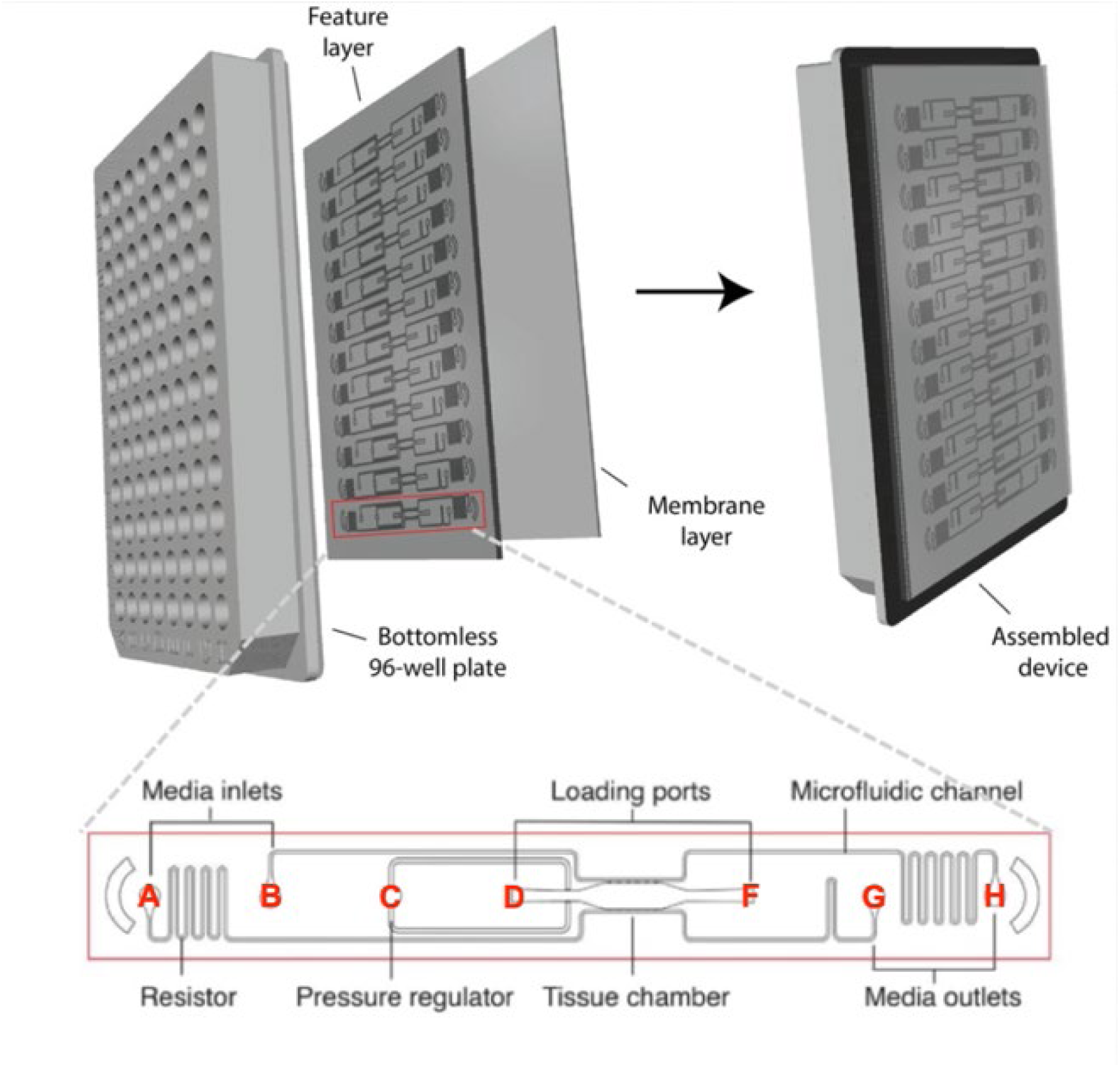
Vascularized micro-organ (VMO) platform. The VMO platform is fabricated with a bottomless 96-well plate plasma-treated to a microfluidic P4 “feature layer” and a membrane layer. One fabricated plate consists of 12 individual VMO devices, each with media inlets and outlets (wells A, B, G, H), loading ports (wells D and F), a pressure regulator outlet (well C), and the tissue chamber.

**Supplementary Figure 2.**
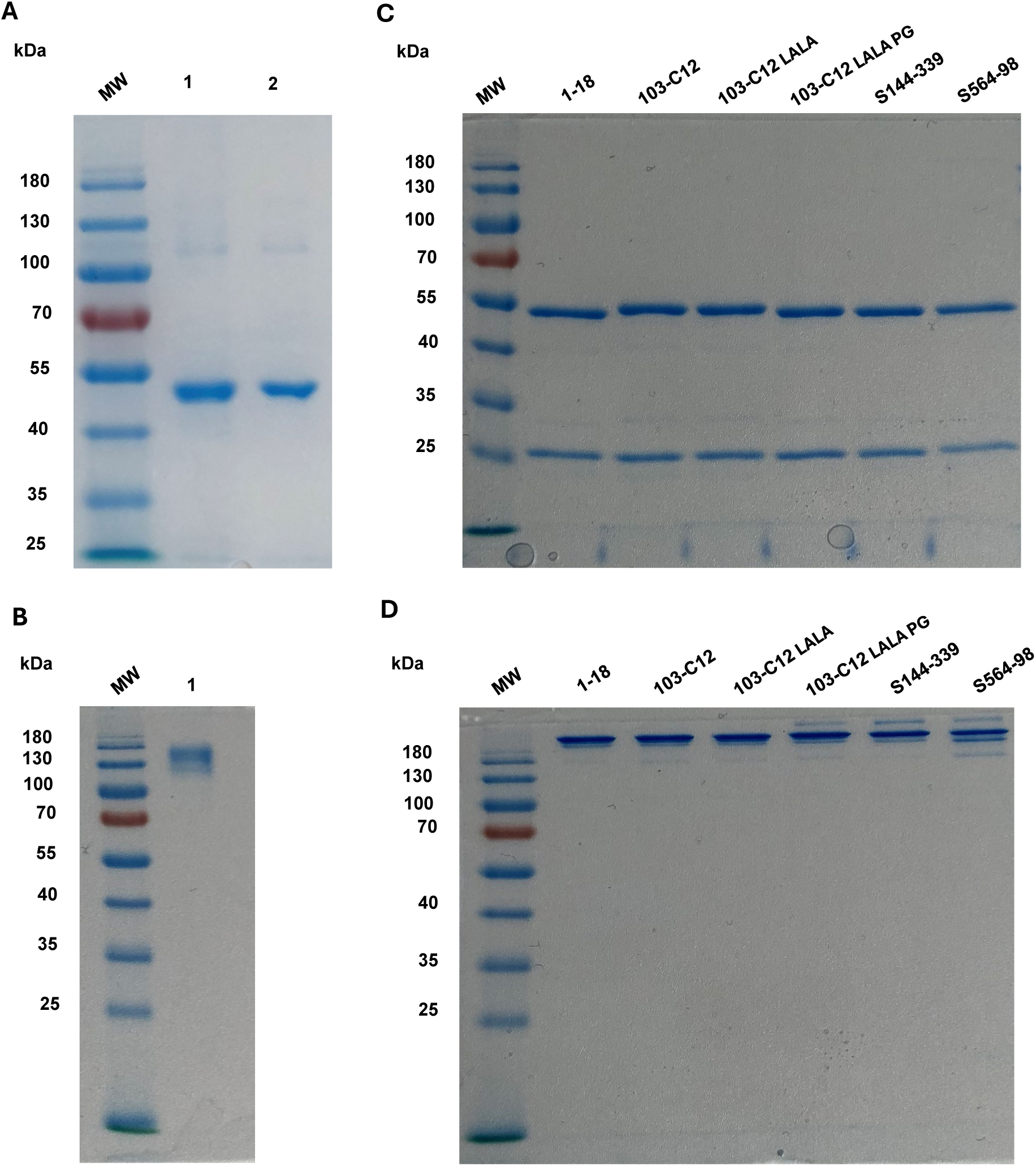
Analysis of SARS-CoV-2 proteins and antibodies. **(A)** Strep-Tactin purified SARS-CoV-2 N (lane 1, ∼47 kDa) displays purity comparable to a commercial standard (lane 2) on a 10% non-reducing SDS-PAGE gel (3 µg loaded/lane). **(B)** Ni-NTA purified recombinant SARS-CoV-2 S (lane 1, ∼130 kDa) runs as a single, highly pure band on a 12% SDS-PAGE gel (2 µg loaded). **(C, D)** Protein A-purified human mAbs targeting SARS-CoV-2 S and N were analyzed under reducing (**C**) and non-reducing (**D**) conditions (1 µg loaded/lane) on a 10% SDS-PAGE gel. Reducing conditions reveal expected heavy (∼50 kDa) and light (∼25 kDa) chains, while non-reducing conditions confirm the presence of intact, assembled antibodies (∼150 kDa), verifying structural integrity across all variants. Proteins were visualized using Simple Blue Safe Stain, and molecular weights were verified with a PageRuler Pre-stained Protein Ladder (MW).

**Supplementary Figure 3.**
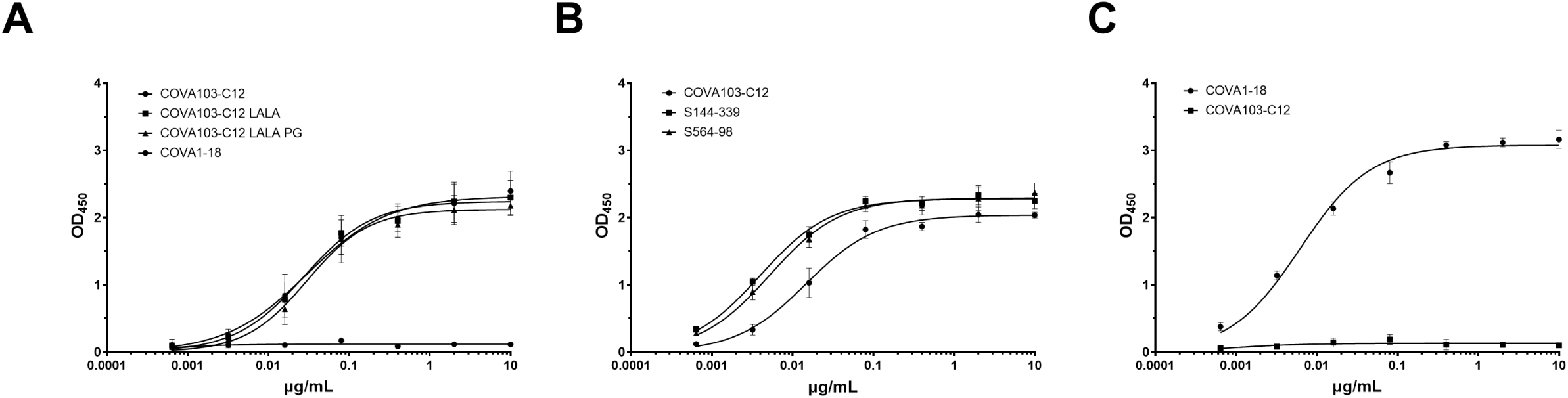
Characterization of antibody specificity and functional integrity. (**A**) Concentration-dependent binding of the parental COVA103-C12 antibody and its Fc-variants, LALA and LALA-PG, demonstrate that high-affinity binding to N is not impacted by Fc-region mutations. (**B**) Comparison of antigen affinities across three N-specific IgG1 antibodies (COVA103-C12, S144-339, and S564-98), with COVA103-C12 exhibiting a slightly lower apparent affinity. (**C**) Specificity confirmation via reciprocal assays, illustrating the high specificity of the S-specific control antibody COVA1-18 for S and lack of binding to S of the N-specific COVA103-C12. Data represent means and standard deviations of ELISAs results repeated 3-4 times with technical replicates.

**Supplementary Figure 4.**
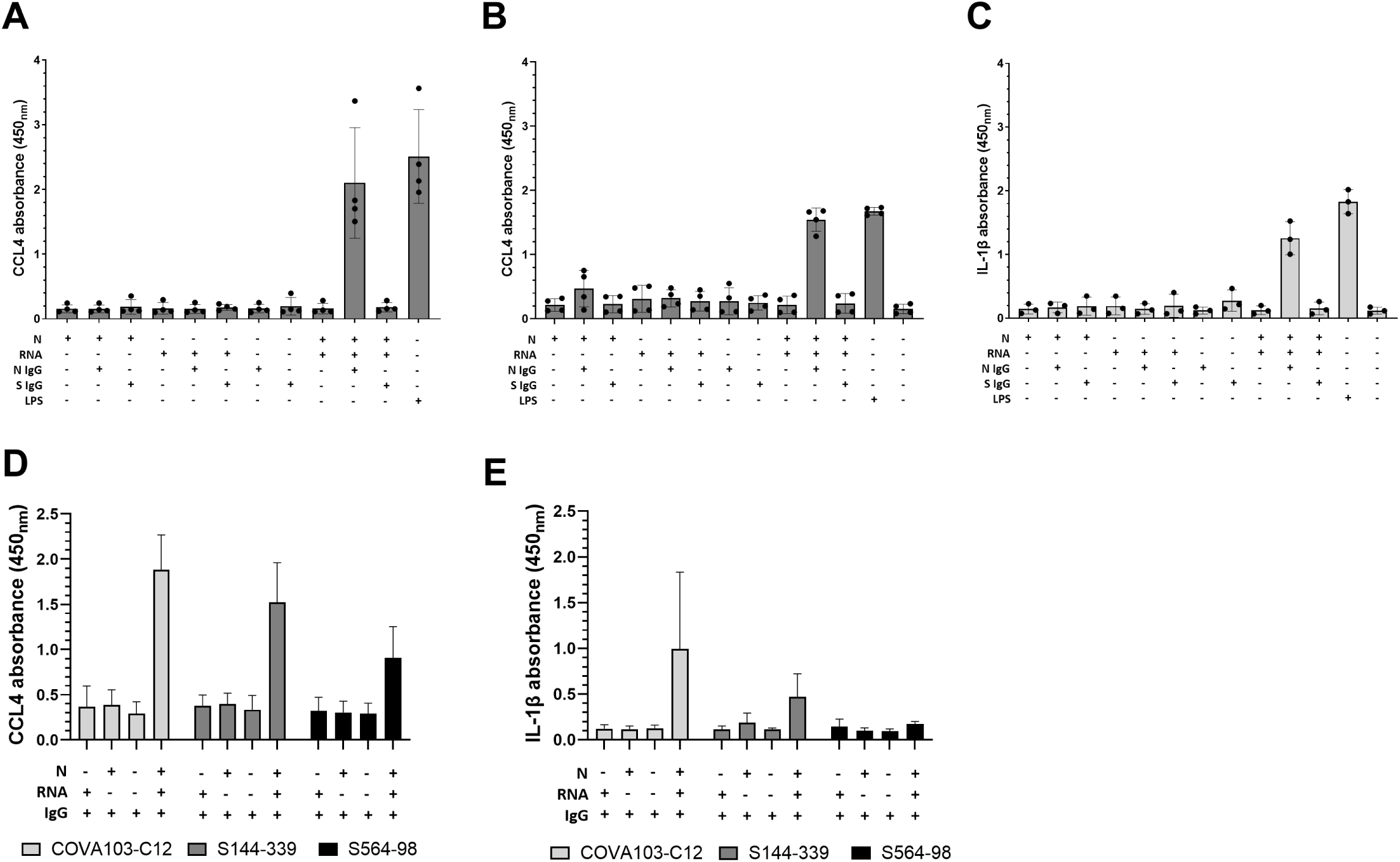
The inflammatory response induced by ICs occurs with different SARS-CoV-2 RNA fragments and with different anti-N IgG1 monoclonal antibodies. ICs were made with a 1.2 kb fragment from the N coding region (**A, D, E**) or a 1.0 kb fragment from the frameshift signal region (**B, C**) of SARS-CoV-2. The antibody component of the ICs consisted of anti-N IgG1 COVA103-C12 (**A**, **B**, **C,**) or of COVA103-C12, S144-339 or S564-98 (**D, E**). CCL4 (**A, B, D**) and IL-1β (**C, E**) were measured in supernatant fluid after stimulating monocytes for 18 hours. Data represent mean and standard deviation of three (**C**) or four (**A**, **B**, **D**, and **E**) independent experiments with different healthy monocyte donors.

**Supplementary Figure 5.**
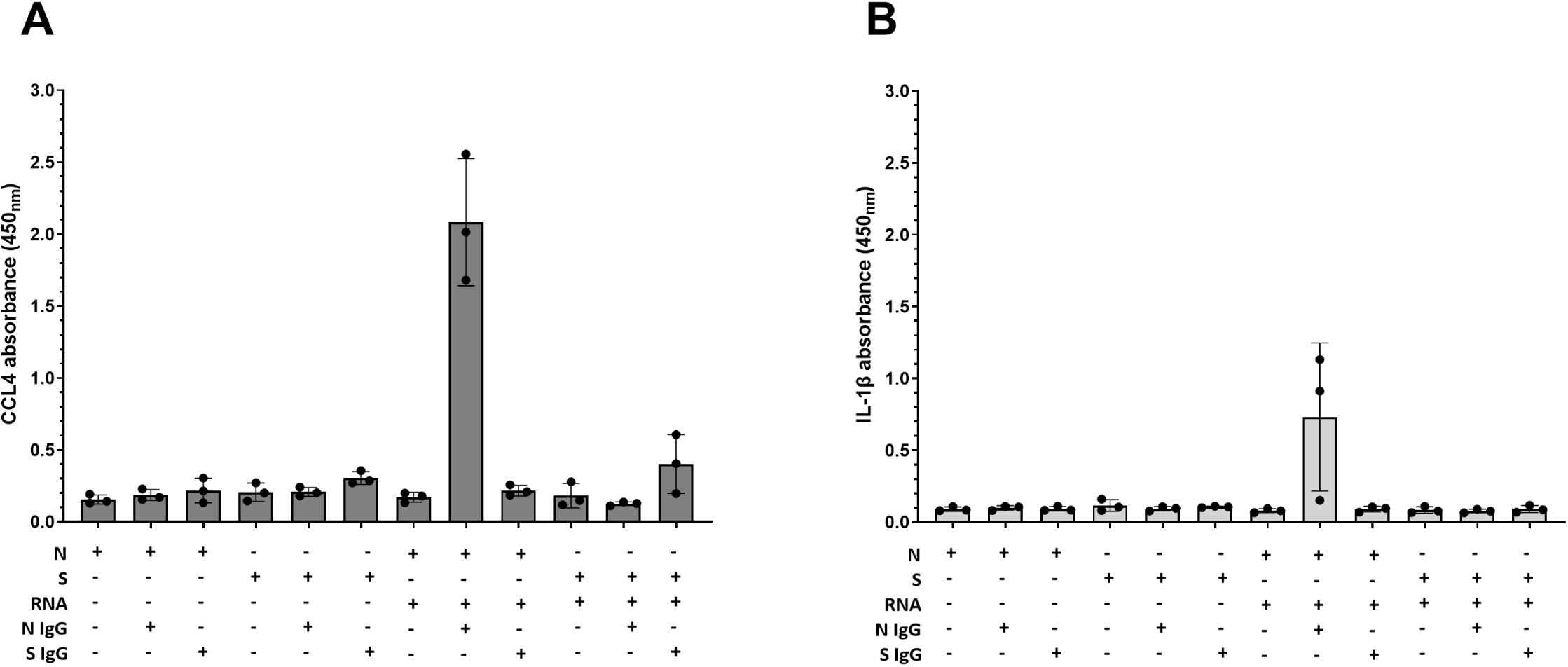
ICs or potential ICs made with S or anti-S IgG1 do not induce an inflammatory response. Primary human monocytes were incubated with ICs or potential ICs made with anti-S or anti-N antibodies, S or N, and RNA. CCL4 (**A**) and IL-1β (**B**) were measured by ELISA in the supernatant fluid of monocytes incubated with the components for 18 hours. Data represent means and standard deviations of three independent experiments with different monocyte donors.

**Supplementary Figure 6.**
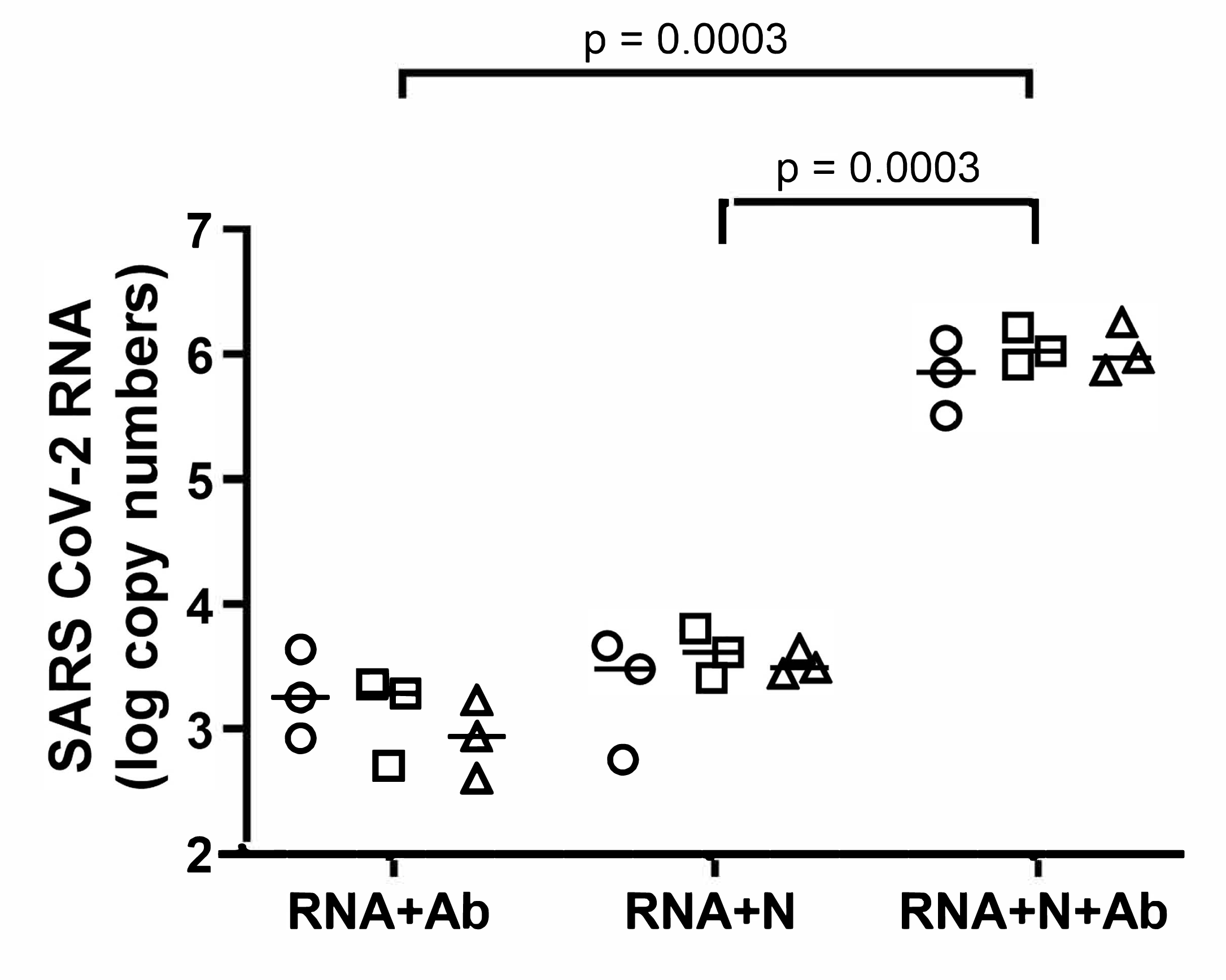
Nucleic acid-containing ICs are internalized by primary human monocytes. Monocytes were incubated with ICs or with controls (RNA+anti-N IgG or RNA+N) for 90 minutes. After careful washing, monocytes were lysed, and the SARS-CoV-2 RNA copies were determined in the lysate by quantitative RT-PCR.

**Supplementary Figure 7.**
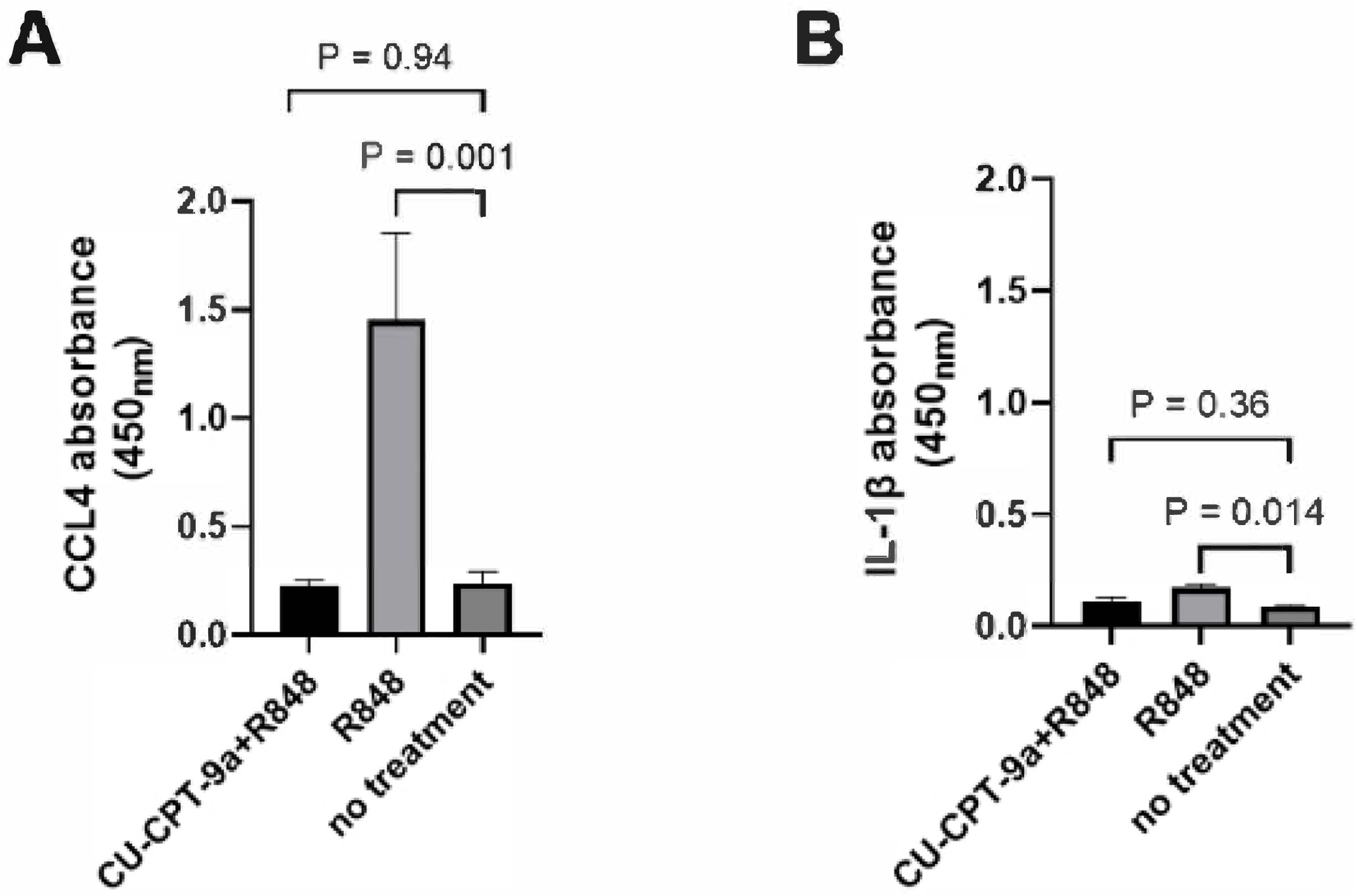
Monocyte TLR7/8 engagement by R848 results in CCL4 and IL-1β production. Data represent means and standard deviations using monocytes from a single donor in triplicate.

